# SynAnno: Interactive Guided Proofreading of Synaptic Annotations

**DOI:** 10.1101/2025.08.09.669342

**Authors:** Leander Lauenburg, Jakob Troidl, Adam Gohain, Zudi Lin, Hanspeter Pfister, Donglai Wei

**Affiliations:** Department of Computer Science, Boston College; School of Engineering & Applied Sciences, Harvard University

**Keywords:** Connectomics, Synaptic Annotations, Neuron-Centric, Proofreading Workflow

## Abstract

Connectomics, a subfield of neuroscience, aims to map and analyze synapse-level wiring diagrams of the nervous system. While recent advances in deep learning have accelerated automated neuron and synapse segmentation, reconstructing accurate connectomes still demands extensive human proofreading to correct segmentation errors. We present SynAnno, an interactive tool designed to streamline and enhance the proofreading of synaptic annotations in large-scale connectomics datasets. SynAnno integrates into existing neuroscience workflows by enabling guided, neuron-centric proofreading. To address the challenges posed by the complex spatial branching of neurons, it introduces a structured workflow with an optimized traversal path and a 3D mini-map for tracking progress. In addition, SynAnno incorporates fine-tuned machine learning models to assist with error detection and correction, reducing the manual burden and increasing proofreading efficiency. We evaluate SynAnno through a user and case study involving seven neuroscience experts. Results show that SynAnno significantly accelerates synapse proofreading while reducing cognitive load and annotation errors through structured guidance and visualization support. The source code and interactive demo are available at: https://github.com/PytorchConnectomics/SynAnno.

## 1 Introduction

Recent advances in connectomics have enabled large-scale reconstruction of neuronal circuits from high-resolution imaging data across diverse model organisms, including Drosophila [10, 27], mouse [39], and even humans [36]. Deep learning-based methods have significantly accelerated this process by automating neuron segmentation and synapse detection. However, despite their high accuracy, these automated approaches still produce errors that necessitate extensive human proofreading [21]. While substantial progress has been made in developing tools and workflows for proofreading neuron segmentation [11, 12], comparatively less attention has been given to the validation and correction of synaptic annotations. This gap poses a critical challenge, as errors in synapse labeling can misrepresent circuit connectivity and lead to biased functional interpretations. As a result, high-quality neuron-centric proofreading of synaptic annotations remains essential for producing interpretable connectomic data.

Neuron-centric proofreading of synaptic annotations is inherently challenging due to the complex branching of neuronal arbors and the sheer volume of synaptic connections in large-scale datasets. For instance, the H01 connectome [36] reconstructs one cubic millimeter of human cortical brain tissue at nanoscale resolution, containing over 57 thousand cells and 150 million synapses-yet only a small subset of its neurons has been fully proofread. Given this immense scale, tools like SynAnno are essential for enabling targeted, high-accuracy annotation of crucial neuronal subsets most relevant for downstream analysis, thereby making large-scale connectome reconstruction efforts more focused and tractable. Traditional proofreading methods typically rely on unstructured manual inspection via 3D visualization tools, a process that is time-consuming and susceptible to inconsistencies.

We present SynAnno, an interactive tool to facilitate guided proof-reading of synaptic annotations in large-scale connectome datasets. SynAnno integrates structured workflows with machine learning-assisted error correction, enabling neuroscientists to systematically review and refine AI-generated synapse masks. The system provides a neuron-centric approach to proofreading, allowing users to traverse synapses associated with each neuron compartment in a structured manner (Fig. 2). To enhance efficiency, SynAnno uses depth-first search (DFS)-based pathfinding for systematic neuron traversal (see Sec. 8), an abstraction pyramid for multi-scale visualization (see Sec. 6.1), and an interactive error-labeling interface (see Sec. 6.1). Additionally, machine learning models help identify and correct synapse mask errors, thereby reducing the overall proofreading burden. We evaluate SynAnno in user and case studies with seven domain experts (see Section 9). We find that our structured proofreading workflow, combined with machine learning-assisted corrections, yields significant improvements in proofreading speed and accuracy. We further discuss the usability insights gained from neuroscience domain experts and highlight opportunities for future improvements. In summary, our contributions are: **(1)** We contribute a guided synapse proofreading workflow. This structured approach allows the effective review of synaptic annotations and incorporates neuron-centric navigation strategies that suggest an optimal proofreading trajectory for each neuron. **(2)** We propose a hierarchical visual interface for error identification and correction. The user first quickly screens groups of synaptic annotations for errors and then further refines the annotations of incorrectly labeled synapses in a separate view. **(3)** We provide an interactive machine learning-assisted error correction approach. SynAnno integrates a deep learning model for interactive synapse mask refinement, improving proofreading efficiency. **(4)** We empirically evaluate SynAnno in a user study with seven neuroscience and proofreading experts, assessing the effectiveness of SynAnno in real-world proofreading tasks.

## 2 Related Work

### Connectomics and Synapse Analysis

Connectomics maps the wiring diagram of brains by identifying neurons and their synaptic connections. A major challenge is analyzing the vast number of synapses in dense circuits. Key objectives include morphology characterization [13, 28], connectivity mapping [10], synaptic classification [14], and synapse density analysis [23]. Despite advances in automated synapse detection [7, 29], challenges remain in ensuring error detection accuracy, identifying false negatives, and correcting erroneous synapse masks and information flow (Fig. 3). Addressing these issues is crucial for generating high-fidelity neural circuit reconstructions.

### Interactive Proofreading for Connectomics

Interactive proofreading tools are vital for correcting errors in automated neuron and synapse segmentations. NeuroBlocks [1] introduced a modular interface for visualizing and editing neural circuits, enabling experts to interactively refine segmentations. FlyWire [11] and NeuTu [45] expanded on this by supporting large-scale, collaborative proofreading of dense connectomic datasets. Haehn et al. [17, 18] proposed guided and semiautomatic workflows that direct user attention to likely segmentation errors for improved efficiency. Later work addressed the scalability of these systems for increasingly large and complex datasets [16]. Tools like VAST [2], VICE [15], and Raveler [9] combine manual and semiautomatic correction with machine learning support, allowing users to iteratively refine annotations. Our work builds on these foundations by introducing a structured, neuron-centric proofreading workflow tailored to synaptic annotations, supporting guided traversal, error categorization, and ML-assisted correction.

### Synaptic Annotation and Focused Proofreading

Synapse annotation at scale remains a major challenge in connectomics. Plaza et al. [32] introduce scalable methods for synapse labeling, while focused proofreading [30] prioritized regions with a high likelihood of annotation errors to improve efficiency. Lin et al. [24] propose an active learning framework that identifies informative synapse instances for manual review, thereby enhancing model performance with minimal human effort. neuPrint [31] offers a user-friendly platform for querying and analyzing synaptic connectivity. Together, these efforts underscore the need for efficient and targeted proofreading strategies to alleviate the manual burden of validating automated annotations.

### Automatic Segmentation Error Correction

Connectomics has employed a range of approaches [8, 20, 42] to address segmentation errors, particularly split and merge mistakes in neuron reconstructions. Zung et al. [47] trained multiscale 3D convolutional networks to detect and correct such errors, while graph-based models [26] represent neurons as annotated graphs to support structured proofreading. Shape correction techniques, including point cloud models [3, 42] and variational autoencoders [38], learn shape priors to recover fragmented or merged segments. Building on these efforts, our method incorporates a 3D U-Net-based synapse segmentation refinement directly into the proofreading workflow, reducing reliance on manual correction and improving efficiency.

### Visualization for Connectomics

Effective visualization is critical for analyzing large-scale connectomics datasets, enabling researchers to navigate and interpret complex neuronal structures [6, 41, 43]. Traditional tools like Neuroglancer [25] provide interactive 3D environments that facilitate the exploration of dense neural reconstructions, yet they can lead to occlusion and cognitive overload. Abstraction-based methods, such as NeuroLines [5], simplify connectivity representations using intuitive metaphors, while ConnectomeExplorer [4] integrates query-based analysis with multi-scale visualization to support comprehensive data exploration. Our approach extends these visualization techniques by incorporating a hierarchical proofreading interface that allows seamless transitions between 2D, quasi-3D, and full 3D views, optimizing the balance between overview and detailed inspection.

## 3 Neuroscience Fundamentals

### Neurons and Synapses

Neurons are the fundamental units of the nervous system, transmitting electrical and chemical signals across the brain. Each neuron comprises a cell body (soma), signal-receiving dendrites, and a signal-sending axon (Fig. 4). Neuronal communication occurs at synapses-specialized junctions where the presynaptic neuron releases neurotransmitters into the synaptic cleft, activating receptors on the postsynaptic neuron. This directional signaling enables the flow of information through neural circuits. Synaptic polarity, determining whether a synapse is excitatory or inhibitory, shapes neuronal computations: excitatory synapses increase, while inhibitory synapses decrease, the likelihood of an action potential.

### Information Flow

The spatial arrangement of synapses is critical for understanding how information flows within neural networks. Synapses are often clustered in specific regions of a neuron, forming functional units known as synaptic boutons on axons and dendritic spines on dendrites. The placement of these synaptic connections determines signal integration and network dynamics.

## 4 Goal & Task Analysis

The development of SynAnno was guided by close collaboration with neuroscience experts who emphasized the need for a more structured, scalable, and efficient approach to proofreading synaptic annotations. These experts required a proofreading workflow that not only systematically guides them through the dataset but also preserves a clear overview of neuron morphology and proofreading progress. At the same time, they need precise control over corrections and the ability to incorporate machine learning-assisted suggestions without compromising accuracy.

Following a design study methodology, we identified a set of domain goals, requirements, and tasks through multiple semi-structured interviews with six neuroscience experts conducted over a two-year period. All experts work on building and analyzing connectomes from Drosophila, mouse, and human datasets and are trained specialists who routinely engage in high-precision synapse proofreading. The interviews were complemented by feedback gathered from three additional experts in informal conversations at connectomics workshops.

### 4.1 Domain Goals

The primary goal of proofreading synaptic annotations in large-scale connectomics datasets is to ensure that reconstructed neural circuits are structurally accurate and biologically interpretable. Given the scale and complexity of current EM datasets, even small annotation errors can propagate into large-scale misinterpretations of connectivity and circuit function. In collaboration with neuroscience experts-including, but not limited to the participants in our user study (see Sec. 9)-and ongoing discussions, we identified five core scientific objectives that define the goals of synapse proofreading:

#### G1 - Ensure Synaptic Validity

Automatically predicted synapses may include false positives due to imaging artifacts, ambiguous morphology, or segmentation errors. A critical goal is to validate that each annotated synapse represents a true biological connection by confirming the presence of structural indicators (e.g., vesicles, clefts, and membrane specializations). Removing invalid predictions helps prevent spurious connections in downstream connectivity analyses.

#### G2 - Increase Synapse Recall

False negatives-real synapses missed by automated models-are common, especially for small or morphologically atypical contacts. Recovering these instances is essential for maintaining the completeness of the reconstructed connectome. Since undetected synapses often occur in complex or low-contrast regions, targeted workflows are necessary to facilitate their discovery.

#### G3 - Guarantee Functional Accuracy

Correctly assigning the directionality and polarity of synapses is critical for interpreting how information flows through a neural circuit. Misannotations-such as incorrect pre- and postsynaptic assignments-can alter the inferred function of a pathway. Proofreading must ensure that synapse annotations reflect biologically plausible connectivity.

#### G4 - Correct Synapse Morphology

Segmentation masks must precisely delineate the extent of each synapse to enable accurate quantification and partner identification. Common issues include merged synapses, fragmented masks, or misaligned boundaries. Improving mask quality through manual or ML-assisted refinement enhances downstream analysis.

#### G5 - Support Reproducibility and Interpretability

Annotations should be consistent and transparent to facilitate collaboration, validation, and cross-dataset comparisons. Clear workflows, structured annotation processes, and visual traceability ensure that results can be reproduced and interpreted by other researchers. Supporting reproducibility also enables integration with computational pipelines for simulation, analysis, and comparative studies.

### 4.2 Design Requirements

To meet the domain goals, we translated key challenges into actionable design requirements, addressing scale, structural complexity, and cognitive load in proofreading large connectomics datasets.

#### R1 - Orientation and Partitioning

Unintuitive partitioning of neural structures can lead to a loss of spatial and structural context during large-scale proofreading, impairing annotators’ awareness of their current location, connectivity to previously reviewed regions, and the neuron’s overall morphology. This weakens their mental model and increases the risk of overlooked sections. The system should therefore provide intuitive visual cues that maintain spatial continuity and clearly associate individual synapses with the broader neuronal structure.

#### R2 - Process Tracking

Poor process tracking can result in redundant work, overlooked areas, and inefficiencies-especially after task interruptions. The system should clearly show review status-completed, current, and remaining-to support effective progress tracking, maintain contextual awareness, and simplify task resumption.

#### R3 – Pathfinding

Determining a coherent traversal path through neuronal structures is difficult due to their extensive size and complex connectivity. Without structured navigation, proofreading risks fragmented coverage and difficulties in task delegation. The system should provide a biologically meaningful, deterministic, and recoverable traversal strategy aligned with neuronal topology, enabling systematic and reliable task management.

#### R4 - Visual Hierarchy

Synapse evaluation complexity varies significantly, e.g., due to orientation in anisotropic volumes, synapse size, and data origin. Many synapses can be rapidly assessed, while others require detailed inspection. The system should implement a visual abstraction hierarchy, defaulting to simplified instance views with easy access to high-resolution details, supporting efficiency and accuracy.

#### R5 - Task Separation

Proofreading involves a sequence of distinct subtasks: detecting potential errors, diagnosing their cause, and applying corrections. Traditional workflows often intermingle these tasks, forcing annotators to switch context frequently. The system should adopt a clear separation-of-concerns approach, enabling annotators to focus on one task at a time and supporting a systematic, efficient, and cognitively manageable workflow.

### 4.3 Tasks

Based on the domain goals and design requirements outlined above, we derived the following key tasks that SynAnno must support:

#### T1 - Classify predicted synapses as correct, incorrect, or unsure

[G1, G5]: Rapidly triage predicted synapse masks using structured navigation (R1, R3), progress tracking (R2), and visual hierarchies (R4) to optimize speed and accuracy while minimizing cognitive load (R5).

#### T2 - Label incorrect or uncertain synapses with specific error types

[G1, G5]: Directly compare problematic instances (R4) and assign detailed error labels (R2) within a distinct workflow step (R5).

#### T3 - Remove predicted synapses that are biologically invalid

[G1, G3]: Identify and delete invalid predictions using detailed inspection (R4), spatial context (R1), and direct comparison (R5).

#### T4 - Correct the direction of information flow for valid synapses

[G3, G5]: Reassign pre- and postsynaptic labels (R4) to accurately represent information flow (R1) within a distinct correction step (R5).

#### T5 - Refine or redraw synapse masks of erroneous synapses

[G4, G5]: Manually adjust or recreate masks with high-resolution inspection (R4) and spatial awareness (R1) to ensure anatomical accuracy (R5).

#### T6 - Discover, add, and annotate missing synapses to the dataset

[G2, G3, G4, G5] Detect false negatives using deterministic traversal (R3), spatial orientation (R1), and visual hierarchies (R4), and add annotations within a focused workflow (R5).

## 5 Neuron-centric Proofreading Workflows

### Overview

Users start by browsing a connectome data set via an image viewer [25] to inspect and select a neuron for proofreading (see Fig. 2a). Next, SynAnno precomputes synapse metadata, downloads the neuron skeleton, and partitions it into anatomical compartments (see Fig. 2b) by generating a depth-first traversal path originating from the soma (see Sec. 8). Finally, SynAnno maps predicted synapses to their respective neuron locations (see Fig. 2c). In the Error Detection view (see Fig. 5ab), SynAnno embeds a 3D skeleton viewer [44] to display the neuron and its synapses (Fig. 5a). Each compartment is color-coded and accessible via a legend (Fig. 1, right). Users are guided through compartments sequentially based on the traversal path, with synapses loaded in the order of the skeleton nodes. Alternatively, users can bypass this sequence by selecting specific compartments directly via the legend. Upon selection, the interface focuses on the corresponding compartment and loads the first page of associated synapses (Fig. 5, left). From here, users can engage in both the *synapse mask correction workflow* and the *false negative correction workflow*.

**Fig. 1:**
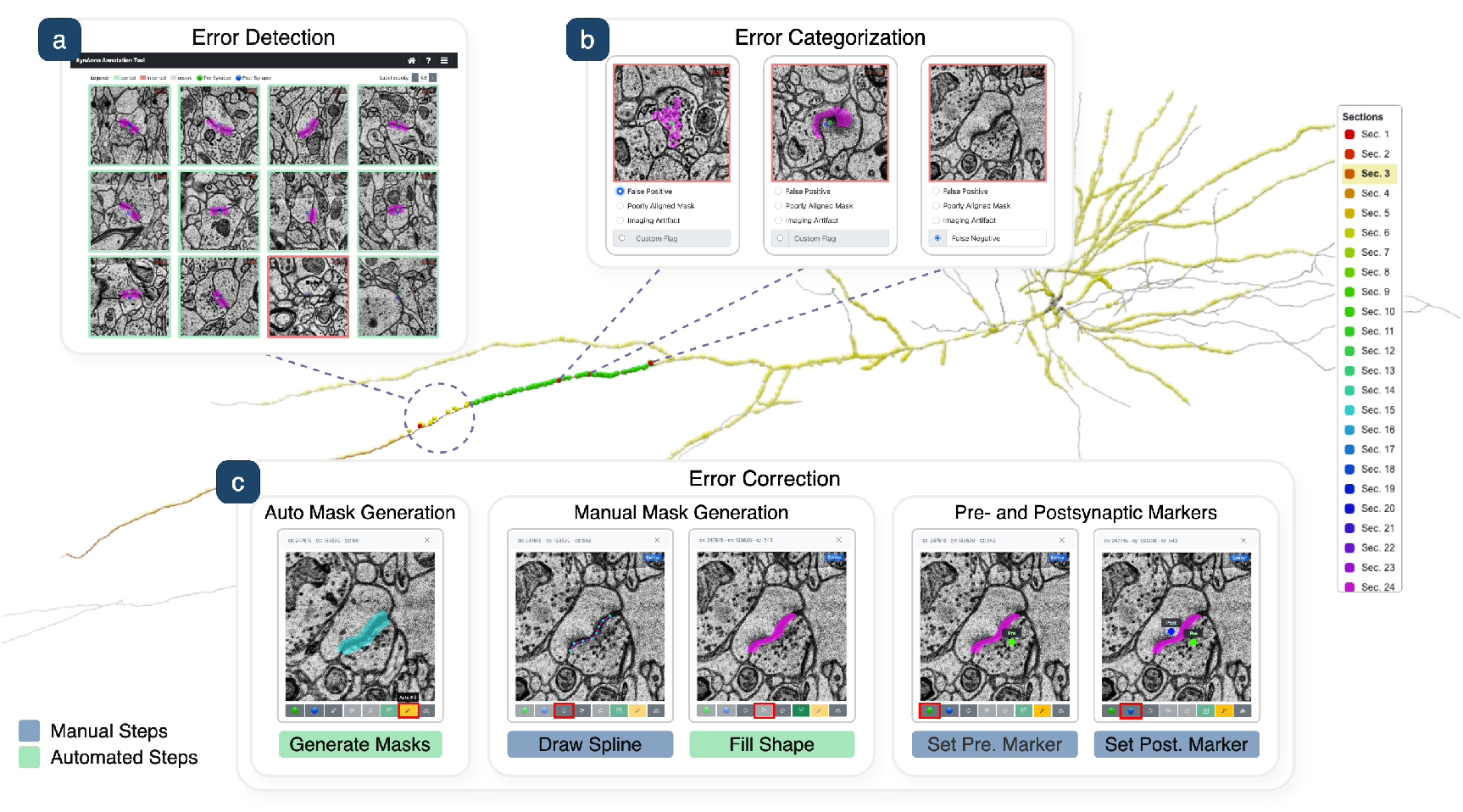
Application Views. SynAnno splits synapse proofreading into three views: (a) Error Detection-users review synapse masks in a grid layout, guided by an interactive 3D neuron viewer; (b) Error Categorization-identified errors are validated and classified; (c) Error Correction-users fix segmentation masks using manual or automatic tools and set pre- and postsynaptic markers.

**Fig. 2:**
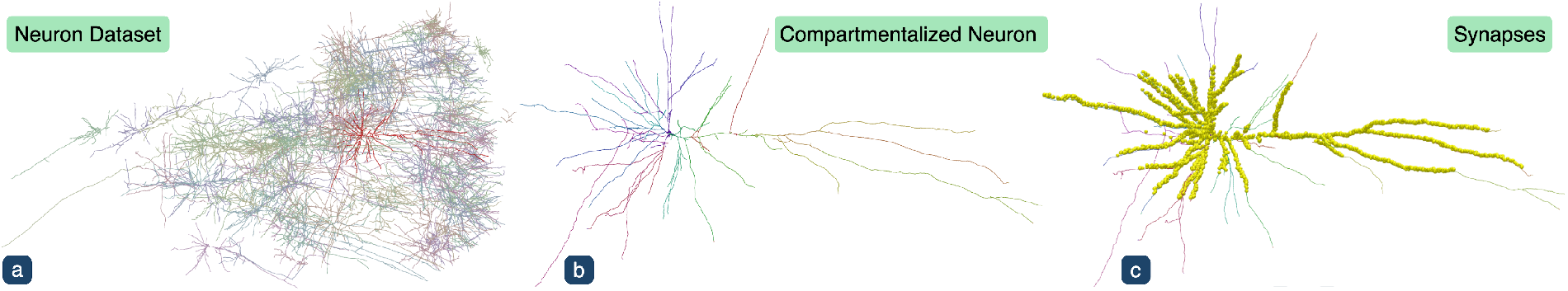
Neuron-Centric Synapse Proofreading. (a) The user selects a specific neuron from the dataset (shown in fully opaque deep red for emphasis). (b) The neuron is compartmentalized into biologically meaningful sections, breaking down the task for focused, systematic proofreading. (c) The user systematically reviews the associated synapses (yellow dots), progressing through the neuron’s compartments.

**Fig. 3:**
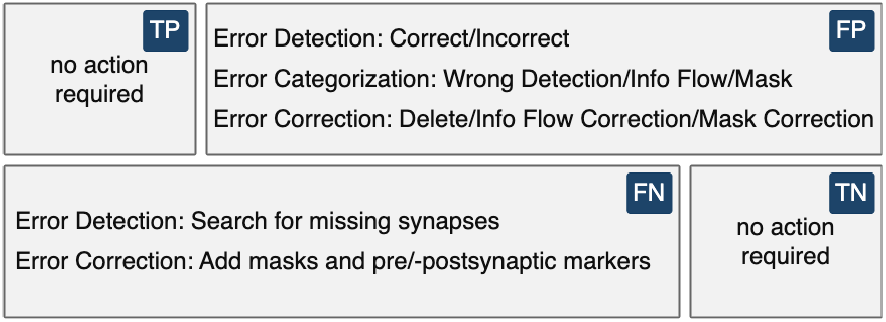
Error Taxonomy. (TP) Correctly annotated synapse. (FP) Errors in detection, information flow, or segmentation require correction. (FN) Requires review of neuron compartments. (TN) No synapse present.

**Fig. 4:**
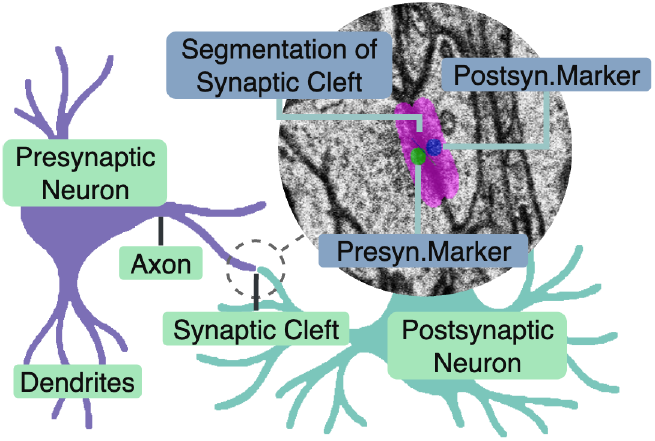
Biological Background. A presynaptic neuron (purple) connects to a postsynaptic neuron (blue) via a synapse. The postsynaptic neuron features a dendritic spine that receives the signal. The inset shows an electron microscopy (EM) image with a segmented synaptic cleft (magenta), presynaptic marker (green), and postsynaptic marker (blue).

**Fig. 5:**
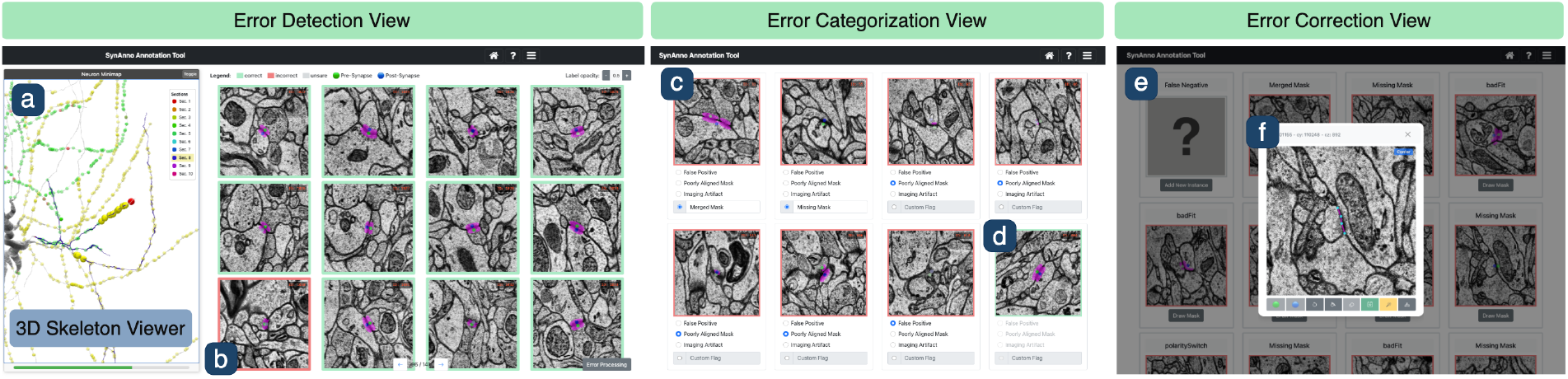
Three Main Views of the Neuron-Centric Workflow. In the *Error Detection* view, users are guided along neuron sections or by the uncertainty-based ordering of synapses within a bounding box. (a) The embedded interactive neuron rendering supports orientation and progress tracking. (b) False-positive synapses can be labeled as incorrect. In the *Error Categorization* view, users revisit instances previously marked as “incorrect” or “unsure” and assign specific error labels, such as (c) “Merged Mask” or (d) reverting the label to “correct.” In the *Error Correction* view, users can (e) add false negatives, (f) manually adjust or auto-generate masks, and reassign pre- and postsynaptic markers.

### 5.1 Synapse Mask Correction Workflow

Figure 6 illustrates the synapse mask correction workflow. Starting with the *Error Detection* view (Fig. 5, left), users review synapse masks and information flow markers, labeling erroneous instances as *incorrect* or *unsure* (Fig. 6a). If additional context is needed, users may inspect the instance using one of three options: an enlarged instance modal view, quasi-3D functionality that enables scrolling through adjacent slices, or a linked image-viewer [25] that directly centers on the synapse mask (Fig. 6b). These optional views help resolve ambiguous cases. Selected instances can be further categorized in the *Error Categorization* view (Fig. 5, center), where users assign predefined error labels or define custom categories (Fig. 6c). Subsequently, users may download the labeled data or proceed to the *Error Correction* view (Fig. 5, right) to correct the synapse mask and adjust the information flow markers (Fig. 6d).

**Fig. 6:**
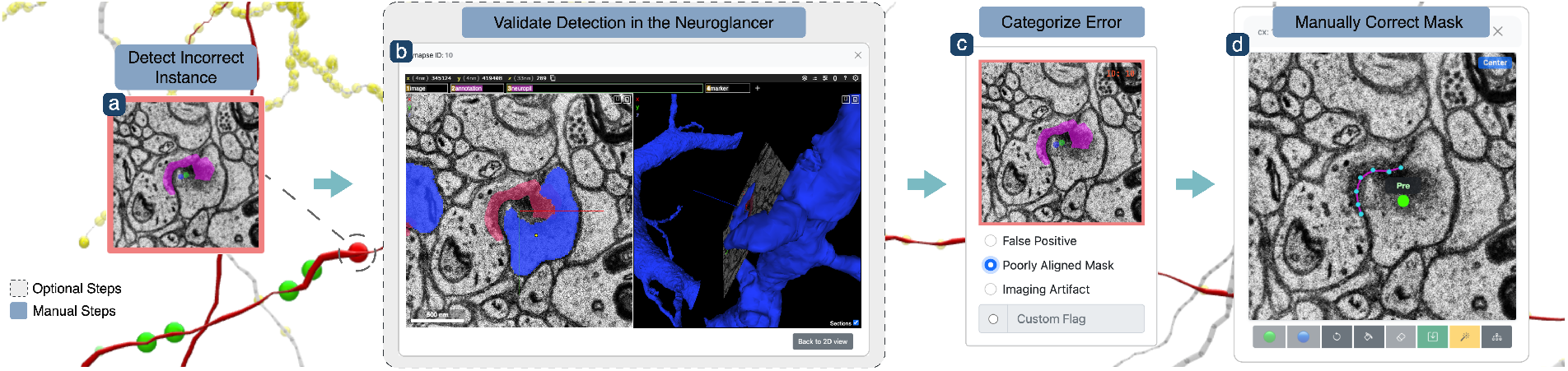
Synapse Mask Correction Workflow. (a) The user reviews a neuron section in the *Error Detection* view, guided by the neuron rendering and identifies a synapse instance they believe is incorrect, marking it accordingly. (b) Optionally, they inspect the instance in Neuroglancer to confirm their suspicion. (c) After completing the review, they assign the label “Poorly Aligned Mask” in the *Error Categorization* view. (d) Finally, they correct the mask and markers indicating the direction of information flow in the *Error Correction* view.

### 5.2 False Negative Correction Workflow

False negatives are systematically searched for during compartment-based proofreading. After a neuron section has been fully reviewed for false positives in the *Error Detection* view, users are prompted to check the same section for missing synapses. Neuron sections can be explored via Neuroglancer, which displays both the neuron skeleton and the surrounding EM context (Fig. 7a). After identifying a candidate synapse, users place a marker at the suspected site. SynAnno then automatically generates a bounding box, which can be adjusted before confirmation (Fig. 7b). Once confirmed, the cropped instance is added to the tile view (Fig. 7c) and labeled as a false negative (Fig. 7d), removing the need for manual categorization. The proofreading process concludes in the *Error Correction* view (Fig. 5, right). SynAnno supports semi-automated segmentation using a 3D U-Net (Sec. 7), with optional manual refinement (Fig. 7e). Additionally, users can place the pre- and postsynaptic markers to indicate the direction of information flow (Fig. 7f), completing the annotation. In addition to neuron-centric proofreading, SynAnno supports a *volume-centric* strategy, allowing users to define a bounding box to review synapses within a specific region, supporting exploratory review or targeted validation.

**Fig. 7:**
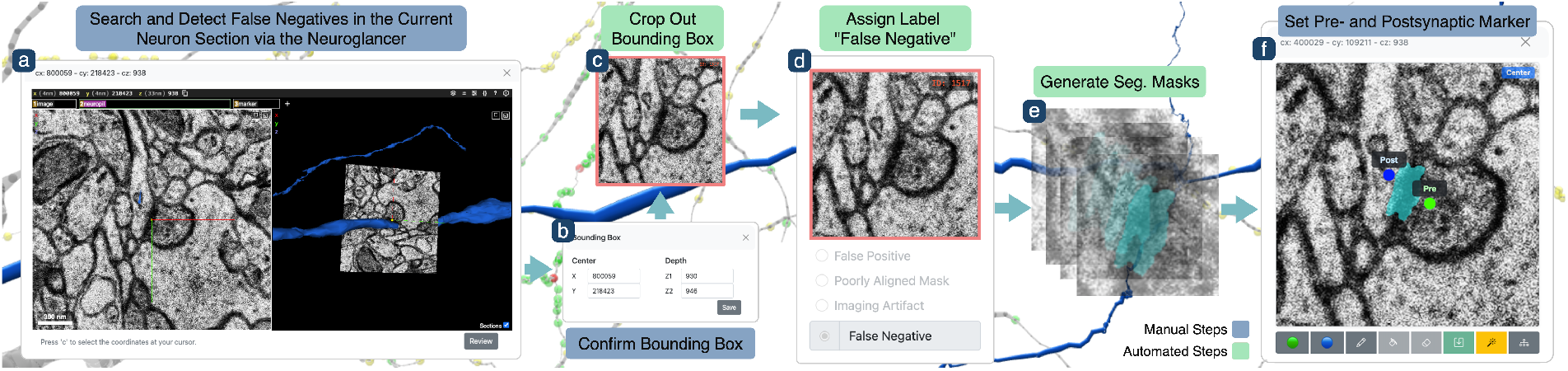
False Negative Correction Workflow. (a) The user navigates along a neuron branch to search for false negatives (FNs) and places a marker at the center of any identified instance. (b) After confirming the bounding box, (c) SynAnno automatically crops the region and (d) labels it as a false negative. (e) In the *Error Correction* view, SynAnno can auto-generate depth-wise synapse masks. (f) The user then sets the markers indicating the direction of the information flow or optionally guides the segmentation process by providing manually drawn masks.

## 6 Interactive Visualization & Interactions

SynAnno provides an interactive visual interface to perform fine-grained synaptic proofreading while maintaining spatial awareness of complex neuronal morphology. The interface supports efficient error detection, clear task separation, and contextualized navigation through large datasets of neurons. SynAnno is organized into three coordinated views: *Error Detection, Error Categorization*, and *Error Correction* (Fig. 5). These views integrate a 2D tile-based instance viewer (Fig. 5b), a 3D slice viewer, a full 3D Neuroglancer integration (Fig. 8), and an interactive 3D skeleton viewer (Fig. 5a), to build a visual hierarchy (R4) and preserve spatial orientation (R1).

**Fig. 8:**
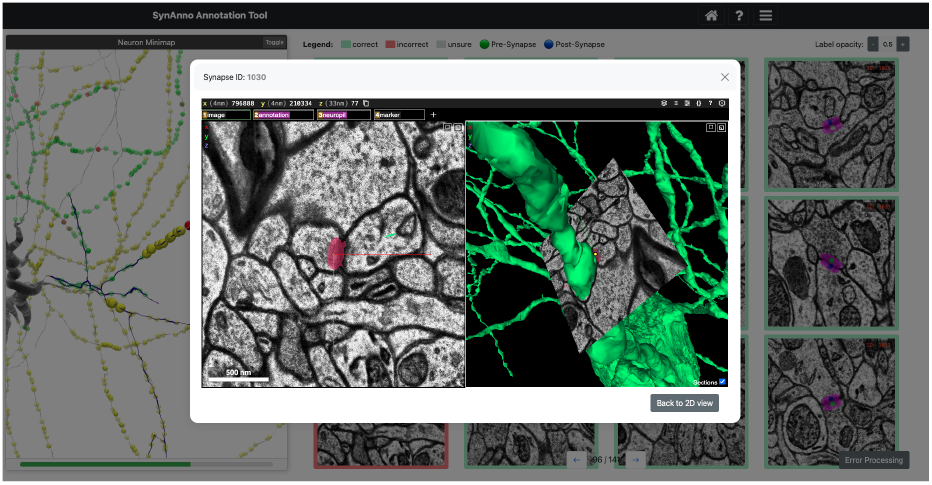
Neuroglancer Integration. In all three main views-*Error Detection, Error Categorization*, and *Error Correction* (Fig. 5)-users can launch a Neuroglancer view, centered on the currently viewed instance. In the *Error Detection* and *Error Correction* views, this enables users to search for and identify false negatives. User study participants highly valued this integration, as emphasized during the semi-structured interviews conducted alongside the usability study and case studies.

**Fig. 9:**
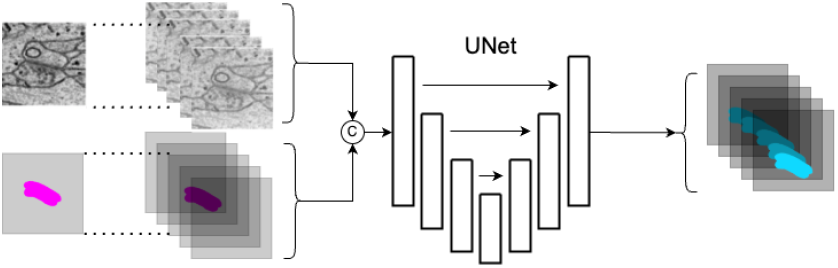
U-Net Model for Synapse Mask Prediction. The electron microscopy (EM) image volume is paired with user-defined seed masks (left, magenta) to form a two-channel input. The corrected masks (right, blue) are auto-generated depthwise around the user’s original mask.

### 6.1 Error Detection View

The Error Detection view addresses the need for scalable triage of predicted synapses (G1, G5) and consists of two parts: (1) a grid of tiles displaying synapses associated with the selected neuron compartment or bounding box (Fig. 5b), and (2) a 3D skeleton viewer visualizing the neuron structure along with its associated synapses. Synapse instances are loaded on a per-page basis; navigating to the next page loads a new subset of instances and updates the 3D viewer accordingly. Users can review each instance and classify it as *correct, incorrect*, or *unsure*, enabling rapid error detection (T1). To minimize manual burden, unlabeled tiles default to the *correct* state when progressing to the next page-streamlining labeling in high-quality regions (R5).

#### Tile-Based Instance Inspection

Synapse instances are rendered as 2D tiles (Fig. 5b), each centered on a representative slice. Users can scroll through adjacent slices directly in the tile to inspect local structures, allowing fast yet informed validation.

#### Visual Hierarchy

For additional structural context (R4), users can utilize an instance-specific module that substantially enlarges the 2D representation, further supporting the examination of adjacent slices. If this level of detail is still insufficient, users can switch to a comprehensive 3D view, which opens at the precise spatial location of the synapse instance. This facilitates multi-scale exploration and reduces ambiguity during evaluation (G1, G3).

#### 3D Skeleton View

To support spatial orientation and reasoning, SynAnno features a fully interactive 3D skeleton viewer (Fig. 5a) that is directly embedded within the Error Detection view. It enables users to rotate, zoom, and pan the currently selected neuron to explore its global morphology and detailed local structure. Synapse locations are visualized as spheres, reflecting their spatial context within the neuron’s morphology (see Sec. 8). The spheres are color-coded by error label-*correct* in green, *incorrect* in red, and *unsure* in yellow. Synapses currently selected in the *Error Detection* view are rendered larger and fully opaque to draw visual attention, while all others appear smaller and semi-transparent to minimize visual clutter. These spatially contextualized visual cues provide real-time feedback, supporting functional accuracy (G3) and structural completeness (G2).

#### Branch Management

To enable consistent proofreading along complex neuron morphologies, SynAnno partitions neurons into multiple compartments, ordered along a depth-first order tree-traversal beginning at the soma (Section 8). This ordering defines a deterministic path, which is visually communicated through compartment-specific colors on the neuron and a legend (R3). Users are prompted to follow this path in the *Error Detection* view. Still, they can also override it through direct interaction with the legend, which enables navigation between sections (Fig. 1, right). Inactive compartments are grayed out, balancing local focus with global context (R1, R2).

#### Visual Synchronization

The 3D skeleton view is tightly integrated with the *Error Detection* view, forming a dynamic, bidirectional connection between spatial and semantic information. Updates made during proofreading are reflected immediately-for example, through a change in the color of a newly annotated synapse-reinforcing a coherent mental model of progress (G5; R2). Synapses currently under review are visually emphasized-enlarged and fully opaque-while others are rendered semi-transparent or downscaled, depending on their review state. As users progress through a compartment by reviewing its synapses in the grid view, the skeleton viewer automatically updates its focus to the currently displayed subset. Once all instances are reviewed, the viewer shifts to the next compartment-determined by a depth-first search (DFS) traversal order (see Section 8)-highlighting the new compartment and dimming the rest, while the grid view begins loading the corresponding synapses. Synapse ordering follows the depth-first traversal of skeleton nodes, ensuring that displayed subsets remain spatially coherent (see Section 8). This automatic progression can be overridden at any time by selecting a compartment from the legend, which loads the first subset of its synapses into the grid and updates the 3D view accordingly. This coordination ensures that users remain oriented and supports structured annotation coverage across views.

#### Progress Tracking

While visual synchronization and branch management provide direct feedback on progress, the viewer also features a global progress bar at the bottom (Fig. 5a). This bar offers an at-aglance summary of proofreading completion, helping users monitor coverage, revisit specific branches, and avoid redundant effort-thereby promoting transparency and sustaining motivation (R2). In addition, users can export and reload their proofreading state to resume or share work across sessions and collaborators (G5) (see Sec. 8).

### 6.2 Error Categorization View

In the *Error Categorization* view (Fig. 5c-d), users can assign predefined error types or define custom labels (T2), directly supporting G1, G3, and G4. This structured stage enforces task separation (R5), enabling users to focus solely on error categorization, and helps maintain annotation transparency and reproducibility (G5). The previously identified synapse instances are displayed in a tile grid, supporting visual hierarchy features. Each tile now includes a set of predefined selectable error categories and an optional field for custom descriptions (see Fig. 1). The two-phase workflow of error detection, followed by categorization, provides a built-in validation loop and enables direct, side-by-side comparison of supposedly incorrect instances. This enables users to amend decisions and restore instances to their correct status, as suggested by participants during the user study (Sec. 9).

#### False Negative Discovery

False negatives-i.e., missing synapses-are identified through interactive exploration in Neuroglancer (Fig. 8), which allows users to flag regions where annotations may be absent (G2). When such a region is marked, SynAnno extracts the local volume and creates a corresponding tile view for synaptic annotation. Users can then place directional markers and generate a new synapse instance (T6). This pipeline supports tasks involving the discovery of false negatives, promoting structural completeness (G2) and functional accuracy (G3), while maintaining spatial context (R1, R3).

### 6.3 Error Correction View

The *Error Correction* view (Fig. 5, right) supports the correction of erroneous or incomplete instances identified during the error categorization. This view addresses the need for anatomical fidelity and functional accuracy (G3, G4) by enabling users to revise synapse masks and reassign directional markers (T4). Instances are presented in a tile grid. Selecting a tile opens the corresponding error correction interface.

#### Marker and Segmentation Correction

SynAnno’s error correction interface (Fig. 5f) enables precise editing of pre- and postsynaptic markers (Fig. 1c, right), which can be interactively placed, adjusted, or removed to ensure the correct direction of information flow (T4; G3). Synapse mask correction (T5) is implemented using a hybrid approach that balances automation and manual control (R4, R5). Corrections can be applied via three mechanisms: *(1)* fully automated using a 3D U-Net, *(2)* semi-automated by guiding the model with user-provided masks on selected slices, or *(3)* fully manual through spline drawing on all individual slices. The manual interface allows users to define masks by placing control points that form filled splines in the style of H01 synapse masks (Fig. 1c, center). An integrated eraser tool supports localized refinement. This strategy accommodates a range of user preferences and preserves anatomical precision (G4; R4, R5). The *Error Correction* view also supports adding and segmenting new synapses (Fig. 5e) that were missed during automated prediction (T6).

## 7 Machine Learning Guided Error Correction

The primary challenge in synapse segmentation lies in accurately identifying each instance and generating an initial 2D mask. Once this first 2D mask was created, annotators must spatially extend the mask to cover the 3D structure of the synapses. This repetitive, slice-by-slice manual annotation compromises efficiency and consistency, particularly when scaling to large datasets. Thus, we implemented a 3D U-Net [34, 48]-based model designed to fully automate the segmentation or volumetrically extend masks across all slices of an instance. Our approach minimizes manual work, allowing annotators to concentrate on higher-level corrections while ensuring consistent segmentation.

### Model Integration

Our 3D U-Net supports users in creating synapse masks for false negatives and instances labeled as *incorrect* in the *Error Detection* view and categorized accordingly in SynAnno’s *Error Categorization* view (Fig. 2a-d). Users can employ the model for fully automated instance segmentation or integrate it into an interactive refinement workflow. There, the user iteratively guides and enhances the model’s output. Rather than relying solely on automated predictions, users can intervene by manually drawing masks on selected slices within the *Error Correction* view (Fig. 2e-f). Once a mask is provided, the model propagates it depthwise by extending the masked region through adjacent slices, using the user-defined mask(s) as a seed to guide continuity and structure. The resulting synapse mask can then be reviewed across slices, with manual and auto-generated masks visually distinguished using different colors for clarity (Fig. 1c, left vs. center). If the output is unsatisfactory, users can refine it by selecting additional key slices, drawing new masks, and re-triggering the synapse segmentation. This adaptive workflow enables flexible intervention, ranging from minimal interventions to fully manual annotation.

### Model Training and Dataset Generation

Our 3D U-Net inputs a dual-channel 3D volume, one channel for the raw EM image and one for a user-defined seed mask, to produce a refined single-channel synapse segmentation. The model was trained on a subset of the H01 dataset, comprising 300 instances for training and 50 for cross-validation. We applied data augmentation techniques, including random flips and rotations, to enhance training diversity. Despite the small training set, the model demonstrated robust performance across a diverse range of scenarios. To emulate realistic user behavior, each instance undergoes multiple segmentation scenarios. One or more seed segmentation layers are retained, while most target slices are hidden from the model. The number of hidden slices is drawn from a normal distribution, introducing variability in the training data. Testing revealed that including instances with zero mask is essential to ensure reliable mask generation in the absence of any seed input and to prevent the model from overcommitting to seed masks.

The model employs a five-level encoder-decoder architecture (32-256 feature channels) and is trained using a weighted binary cross-entropy loss between predicted masks and ground truth masks. Optimization is performed with the Adam optimizer [22], and model selection is based on validation loss checkpointing. Training proceeds for up to 400 epochs with early stopping (patience of 25) and an initial learning rate of 5*×*10^*−*5^, reduced on plateau by a factor of 0.5. All experiments were conducted on a SLURM-managed compute node with 1×NVIDIA A10 GPU, 4 CPU cores, and 128 GB of RAM.

## 8 Data & Implementation

SynAnno relies on five core data sources: (1) raw EM images, (2) synapse masks, (3) neuron masks, (4) neuron skeletons, and (5) meta-data such as voxel size. These sources collectively provide the structural context needed for guided proofreading of synaptic connectivity. Neuron segmentation supports the selection and isolation of individual neurons for focused analysis. Skeletons provide a simplified geometric representation for rendering, compartmentalization, and pathfinding, enabling users to navigate and interpret neuronal morphology.

### Neuron Skeletons

We obtain the neuron skeletons via H01’s cloud storage using the CloudVolume [37] library. Spatial coordinates were normalized to nanometer units. To facilitate downstream partitioning, we pruned peripheral branches that did not significantly contribute to the neuron’s structural or functional interpretation. Upon user selection, SynAnno retrieves the corresponding neuron skeleton and partitions it into structurally coherent segments. (Fig. 2b). Each neuron is modeled as an undirected graph, rooted at the soma when available; otherwise, a central node is estimated using the PageRank algorithm. A DFS traversal imposes a hierarchical node ordering based on traversal indices. Branch points-nodes with degree ≥3-serve as natural mask boundaries. To avoid over-fragmentation, small segments are merged with adjacent ones based on traversal continuity and connectivity constraints. The resulting segments are then ordered to reflect biologically meaningful hierarchies, enabling efficient, compartment-wise proofreading. Skeleton processing, partitioning, and pathfinding were implemented using the Navis [35] and NetworkX [19] libraries. For the usability studies, case studies, and the interactive demo, we curated a subset of 104 morphologically complete neurons.

### Synapse Data

Synapse masks and metadata were retrieved from the H01 cloud storage and the associated connections database. Each instance is defined by a mask highlighting the synaptic cleft and a set of coordinates for the pre- and postsynaptic partners (see Figure 4). To establish spatial correspondence between neuronal morphology and synaptic connectivity, synapse coordinates were extracted and converted from voxel space to nanometers, forming a spatial point cloud. The synapse table was filtered to retain only synapses associated with the selected neurons. A KDTree was constructed from the skeleton node coordinates to enable efficient nearest-neighbor queries. Each synapse was mapped to its closest skeleton node, with a small random offset applied after pruning to prevent overlaps, and its corresponding structural compartment and traversal index recorded. To support structured proofreading within each compartment, synapses were sorted by their assigned compartment and skeleton node’s traversal index.

### Collaboration and Reproducibility

SynAnno’s neuron-centric design partitions the dataset into biologically meaningful compartments, enabling intuitive task delegation within distributed proofreading workflows. This compartmentalization enables users to focus on manageable units, supporting the collaborative and scalable reconstruction of large-scale connectomes. SynAnno also lays the foundation for resuming and sharing proofreading sessions. All user actions-such as labeling and correcting synapses-are automatically recorded. Users can export a JSON file that captures the current proofreading state and extends the materialization table with annotation metadata. This file can be reloaded to resume work or shared with collaborators to continue or review annotations. Additionally, corrected synapse masks can be downloaded, with filenames that encode spatial coordinates to support seamless patch integration into the target volume via CloudVolume. While the current prototype supports only single-user sessions, it has been designed with scalability in mind. A cloud-based infrastructure could readily extend this functionality by synchronizing session data across users, supporting real-time collaboration.

### System

SynAnno is implemented using Python’s Flask framework and vanilla JavaScript. The open-source code and a demo are available: https://github.com/PytorchConnectomics/SynAnno.

## 9 Evaluation

We conducted a qualitative user study with seven experienced connectome proofreaders, a post-session survey, and a set of in-depth case studies targeting critical annotation and proofreading tasks. As participants engaged with SynAnno, we asked structured and open-ended questions to gather task-specific insights and spontaneous feedback. Sessions were video-recorded, transcribed, and thematically analyzed to identify recurring usability challenges and guide iterative refinement. The study involved two postdoctoral neuroscientists, who ensured biological validity, and five professional proofreaders from a leading U.S. neuroscience center, representing the tool’s primary user base. All participants are actively using established platforms such as FlyWire [11], VAST [2], and Neuroglancer [25].

The evaluation was designed to address three core questions: **(1)** How well does *SynAnno* support the cognitive workflows of expert annotators? **(2)** Which aspects of the interface and interaction design facilitate or hinder efficient proofreading? **(3)** What improvements could enhance user experience and task performance? We first present the results of the user study, followed by detailed case studies and a summary of updates implemented in direct response to user feedback.

### 9.1 User Study

We conducted seven semi-structured interviews with domain experts to evaluate the usability and effectiveness of SynAnno. Each session lasted approximately one hour. Following a brief introduction to SynAnno, participants were invited to independently steer the interface and test the workflows based on their expectations and annotation practices. A short follow-up survey was distributed after the session.

#### Survey Results

The survey included eight questions rated on a five-point Likert scale (Strongly Disagree, Disagree, Neutral, Agree, Strongly Agree), covering key aspects such as workflow support, navigation, segmentation, and labeling. Figure 10 summarizes the aggregated survey responses. Each row represents a specific question (Q1-Q8). Neutral responses are centered around the y-axis, with disagreement extending to the left and agreement to the right. Percentages on the right side of each bar indicate the share of users who agreed with each statement. The three highest-rated items-Q1 (workflow support), Q2 (intuitive labeling), and Q3 (dataset navigation)-each received 100% agreement, highlighting these as key strengths of the tool. Statements on general proofreading efficiency and ML-assisted synapse segmentation (Q4-Q5) received 85% agreement. Slightly lower scores were recorded for Q6 and Q7 (both 71%), which addressed logical neuron traversal and progress tracking, respectively. Q8 (false negative handling) received the lowest score (57%). Feedback on Q4 suggests that some participants, particularly those involved in initial synapse segmentation or region-of-interest identification, were less familiar with workflows for correcting existing masks. Concerning Q5, the discussion focused on when synapse masks are necessary. For workflows focused solely on annotating the direction of information flow, synapse mask correction was deemed less relevant. However, for tasks such as estimating contact areas, the ML-assisted synapse segmentation was considered highly beneficial. Responses to Q5 and Q6 aligned with suggestions for structural improvements, including synapse-weighted partitioning, manual compartment definition, and hierarchical skeleton representations in the legend. Q8 reflects concerns voiced during interviews regarding the reliable identification of false negatives. Participants emphasized the need to traverse entire compartments and noted a cognitive disconnect between proofreading and initial annotation tasks. Participants’ qualitative feedbacks are summarized below.

**Fig. 10:**
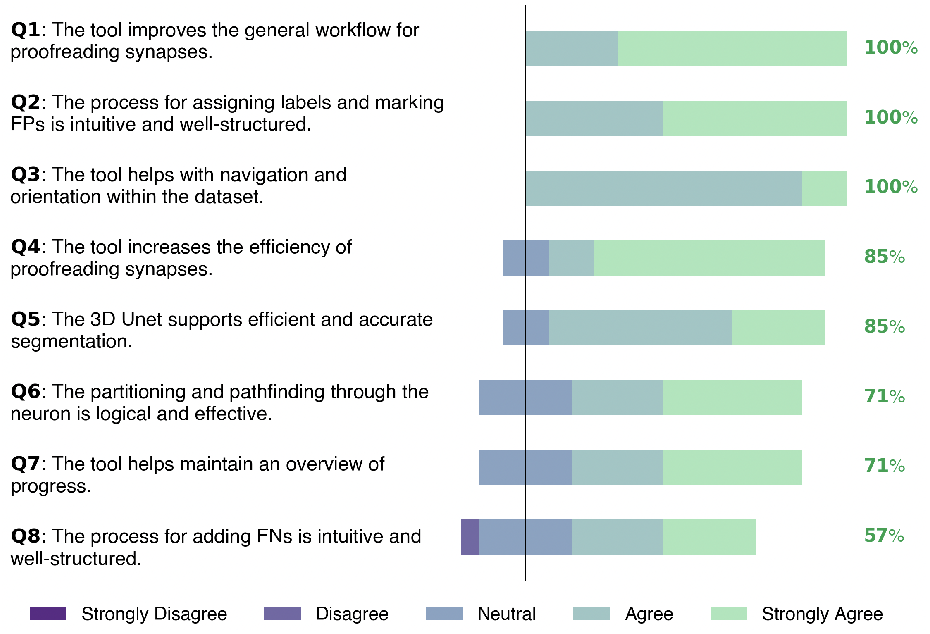
User Study Results. We collected aggregated survey responses from *N*=7 users. The results show strong satisfaction with core functionalities, including workflow support, intuitive labeling, and navigation (Q1-Q3), each receiving 100% agreement. Moderate agreement was observed for ML-assisted segmentation and proofreading progress tracking (Q4-Q7). The lowest score was given to the intuitiveness of adding false negatives (Q8), indicating a need for improvement.

#### Interface and Visualization

A clear point of consensus was the need for larger, resizable tiles in the *Error Detection* view (Fig. 5, left), along with the dynamic adjustment of tile size and slice count based on dataset properties and available screen space. The integration of Neuroglancer (Fig. 8) was widely praised as a powerful feature for context-rich exploration of challenging instances. One participant suggested enabling direct access to Neuroglancer to streamline context switching. Others, in turn, proposed further enhancing the instance module view to eliminate the need to switch to Neuroglancer entirely. Several usability enhancements were suggested to improve the overall interface, such as keyboard shortcut-based navigation and making neuron branches selectable via a clickable skeleton in the minimap. The contextual visual cues provided by skeleton rendering (Fig. 5a) were frequently cited as valuable for supporting orientation.

#### Workflow and Task Logic

A well-received feature of SynAnno was its structured, section-based review workflow (Fig. 6). Participants consistently reported that this approach supported sustained cognitive focus and enabled systematic traversal of the neuron. The ability to pause and resume work at the exact location was described as a transformative capability for long-term annotation projects. Participants expressed divergent views on task flow structure. Some preferred immediate labeling after inspecting an instance to avoid the cognitive burden of revisiting items. Others favored separating review from labeling, emphasizing benefits such as self-checking, comparative validation, and more accurate error correction. Progress tracking was consistently highlighted as a key feature. Individual participants requested additional visual aids, including section-level progress bars and markers for reviewed instances in the tile view to reduce duplicate work. Improved partitioning strategies were also discussed. One key recommendation was to balance the number of synapses per section to promote equitable workload distribution and allow users to manually partition neurons, offering greater flexibility in managing complex data regions.

#### Error Handling and Labeling

Participants offered nuanced feed-back on error labeling strategies, emphasizing the need for additional predefined labels such as *Missing Mask* and *Merged Mask* to enhance error detection precision and filtering efficiency while reducing reliance on manually created custom labels. A significant request was the ability to define custom error vocabularies on a per-dataset basis to accommodate domain-specific terminology and validation objectives.

One of the most debated topics was including an **“unsure”** label. Several participants regarded this option as essential for managing ambiguous cases, enabling deferred judgment or later comparison in the *Error Categorization* view. They expressed concern that forcing a binary choice between **“correct”** and **“incorrect”** could introduce decision bias. Conversely, others worried that the **“unsure”** label could reduce accountability, encouraging annotators to overuse it to avoid making difficult decisions. The prevailing recommendation was to make the availability of this label configurable, allowing individual labs to tailor its use according to their review policies. The *Error Categorization* view was widely praised for supporting iterative proofreading. However, alternative modes were also proposed, such as a swipe-based interface, which would present one instance at a time.

#### False Negative Detection

False negative (FN) detection emerged as one of the most challenging and contested aspects of the annotation workflow. Participants consistently emphasized that accurately identifying FNs requires systematic, branch-by-branch traversal of the entire neuron. This makes the process not only cognitively demanding but also difficult to integrate seamlessly into standard instance-based review flows. Although the idea of embedding FN detection directly into the main review interface was met with appreciation, it became clear that this approach is quite different from the synapse mask review. Attempting to combine both so closely led to the loss of focus, increased error risk, and a breakdown in the logical flow of work. Therefore, some participants advocated for explicitly separating FN detection into its dedicated workflow phase, allowing users to adopt a different mindset and strategy for this task. This separation would also enable optimizations such as tailored tools for scanning long branches, toggling neuron skeleton visibility, and navigating via section or projection views.

#### Synapse Masks Correction and Tooling

Users appreciated the ability to directly correct synapse masks within SynAnno, highlighting the marker correction feature-used to indicate the direction of information flow-for its speed and efficiency. The option to place markers on arbitrary, independent slices was well-received for its flexibility. The U-Net autocomplete feature was a major strength, reducing redundancy and fatigue during review. However, users expressed a need for more manual control, suggesting features like a brush tool for mask painting, adjustable auto-segmentation, and finer control over mask radius.

### 9.2 Case Study

We conducted four case studies **(C1-C4)**, each involving a single expert.

#### C1 - Detecting Erroneous Synapses

Here, the user reviewed pre-existing synapse masks using the *Error Detection* View (Fig. 5b), labeling each instance as *correct, incorrect*, or *unsure*. They noted that the grid-based layout enabled fast and structured progression through the neuron, unlike their usual workflow of manually navigating 3D volumes using spreadsheets. However, the small, fixed tile size was seen as a bottleneck, as it required frequent context switching. Thus, we made the number of tiles per page configurable to accommodate different screen sizes and user preferences. While the user praised the structured flow and the ability to progress quickly through the neuron, they may have overused the *unsure* label to delegate complex cases for supposed later review, reinforcing earlier discussions around annotation confidence and reviewer accountability. The user also requested a feature to track which instances had already been reviewed on a page. At the time of testing, 24 instances were shown per page without any additional identifiers; we have since added visible instance IDs in the top-right corner of each tile to address this need.

#### C2 - Identifying False Positive Synapses

Here, the user was asked to review the synapse masks of five neuron compartments using the *Error Categorization* View (Fig. 5c) to identify false positive synaptic annotations. The task was completed with surprising speed and confidence, although the user relied heavily on Neuroglancer. Compared to a previous dataset of a fly brain, the H01 dataset was perceived as more challenging, with synapses exhibiting less pronounced features. Consequently, the user suggested adding an option to bypass the 3D skeleton view and jump directly into the more detailed 3D viewer, reducing the number of context switches. Overall, the user quickly adapted to the tool and inquired when it would be available in their daily workflow.

#### C3 - Correcting Synapse Masks

This task involved correcting poorly aligned or incomplete synapse masks using the drawing module and U-Net-assisted auto-completion using the *Error Correction* View (Fig. 5d). The user was asked to correct the masks slice by slice manually. For a second subset, we enabled support for the 3D U-Net, allowing both fully automated and semi-automated synapse segmentation. The U-Net was highly praised for enabling users to select an arbitrary slice as seed inputs. Overall, the tool significantly reduced the repetitiveness of mask drawing. The user observed that drawing a single representative slice could dramatically improve the quality of the synapse mask. However, they requested a tool to make corrections to auto-generated masks, like freeform adjustments.

#### C4 - Searching and Adding False Negative Synapses

This task combined exploratory traversal with instance creation, including marking the centers of false negative (FN) candidates, assigning markers that indicate the direction of information flow, and generating synapse masks. The user navigated to a selected neuron section, identified unannotated synapses, and marked their center points. SynAnno extracted the regions, automatically labeled them as FNs, and enabled rapid annotation. The U-Net auto-completion and marker placement tools were seen as user-friendly and efficient (Fig. 7). Nonetheless, this case study confirmed that false negative detection remains the most cognitively demanding task. Users requested visual aids-such as heatmaps or section-level progress indicators-to help avoid missed regions. The session also sparked a discussion on whether FN detection should be integrated into the proofreading interface or moved to a dedicated view. Overall, FN detection remains the most persistent challenge.

### 9.3 Incorporating User Feedback

Based on insights from the recordings of the usability and case studies, we implemented several improvements. One notable enhancement is the revised drawing module (Fig. 11). We increased the image size, removed the slider, and introduced mouse wheel-based slice navigation. The slider was initially included to provide precise control over the displayed slice. However, we replaced it with wheel-based scrolling to reduce interaction complexity and align the user experience with that of the *Error Detection* and *Error Correction* Views. To address orientation concerns, we introduced a tag marking the center slice in the top-left corner. Additionally, slice coordinates are now displayed and dynamically updated in the header as users scroll, helping them maintain spatial awareness. To streamline interactions further, we implemented keyboard shortcuts for essential actions such as closing the window and re-adding pre- or postsynaptic markers. The enlarged image and module layout are designed to decrease the need to open Neuroglancer, while also aligning with user preferences for a more self-contained workflow. In response to user feedback, we also improved the visibility of splines by enhancing their highlighting prior to color filling. Further improvements include the ability to revise annotations directly in the *Error Categorization* view (Fig. 5d). The number of displayed tiles can now be configured based on dataset complexity, screen size, and user preference. Instance IDs were added to each tile in the *Error Detection* view (Fig. 5, left) to help users better track and distinguish them, showing them directly on the image tiles. Additionally, navigation has been improved: users can now navigate directly to neuron sections, rather than having to page through the interface to reach a particular section. Finally, we followed up with the relevant participants from the usability study, identified through our interview transcripts, to validate these changes. The participants confirmed that the enhancements closely aligned with their feedback and significantly improved the workflow.

**Fig. 11:**
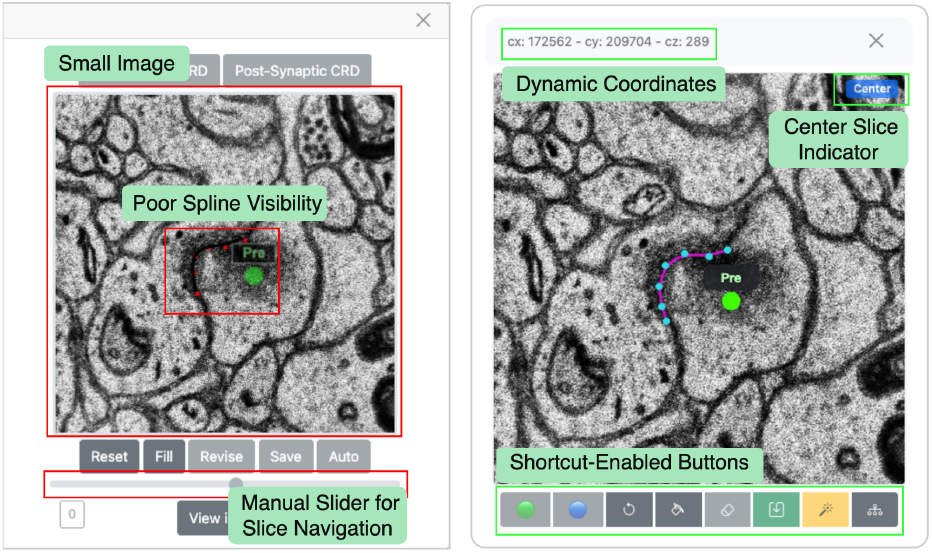
Refactored Drawing Module. Updated drawing module based on user feedback, featuring mouse wheel image slice scrolling, a center slice indicator, dynamic coordinates, keyboard shortcuts, and improved spline highlighting. The figure depicts the initial version to the right and the refactored version to the left.

## 10 Discussion

### User Expertise in Synapse Proofreading

Identifying experts for our user study was challenging, as large-scale synapse proofreading is a highly specialized task. Although the availability of high-resolution connectomics datasets has grown, few labs perform extensive manual proofreading because verifying AI-generated synapse masks remains tedious. By introducing SynAnno, we aim to lower the barrier to connectomics proofreading and encourage broader tool adoption.

### Limitations

SynAnno is designed for synaptic annotation and structured proofreading of neuron-specific synapse collections. It does not support full connectome-wide comparisons of synaptic patterns or large-scale statistical analysis of synapse distributions. While our 3D U-Net model aids in synapse mask correction, its performance depends on the quality of the training data. Challenging cases, such as synapses near dense axonal crossings, may still require substantial manual correction. Additionally, the instance selection algorithm may fail to resolve overlapping synapse masks, requiring manual distinction of adjacent instances. False negative detection remains cognitively demanding, indicating the need for additional visual aids or dedicated interfaces.

### Cross-Domain Applications

SynAnno’s core design offers a generalizable methodology for diverse spatial biology challenges. Its interactive, neuron-centric framework is highly adaptable for other annotation tasks, such as large-scale mitochondria mapping efforts [33]. Furthermore, its fusion of multi-scale visualization and machine learning-assisted correction addresses the analytical needs of emerging fields like spatial transcriptomics, where preserving spatial context and ensuring data quality are paramount [40, 46]. Consequently, our design makes SynAnno a valuable tool for proofreading high-resolution, spatially organized biological datasets.

### Balancing Automation and User Control

Automated synapse segmentation reduces manual annotation effort, but excessive automation can introduce systematic errors and diminish user engagement in critical decisions. Our approach enables users to interactively validate and correct mistakes with AI-assisted guidance while preserving user control. Future versions of SynAnno will offer more adaptive controls, such as manual neuron partitioning and synapse refinement.

## 11 Conclusion and Future Work

By combining a neuron-centric workflow, hierarchical visualization, and machine learning-assisted error correction, SynAnno enables efficient validation and correction of synapse masks for accurate neuronal connectivity reconstructions. SynAnno’s structured, multi-view workflow balances high-throughput triage with fine-grained editing-principles relevant to any visual analytics task involving large, spatially organized datasets. Integrating semantic task separation, hierarchical abstraction, and biologically informed path traversal provides a blueprint for scalable, efficient interfaces, offering insights for interactive visualization systems in fields such as medical imaging, spatial omics, and digital pathology. Looking ahead, we plan to expand 3D U-Net training and implement online learning that adapts to challenging cases by incorporating manually corrected instances from the *Error Correction* view. Furthermore, we aspire to integrate SynAnno into existing ecosystems and workflows to enhance its scalability and support collaborative use. Additionally, we aim to extend SynAnno ‘s applicability to other neuronal structures, further broadening its impact. Enhancing user-driven neuron partitioning and enabling manual coloring and sorting of compartments will offer greater flexibility for complex architectures. Refining our instance selection algorithm will improve segmentation handling, reducing annotation overlap. As connectomes grow in scale, SynAnno will evolve to meet the demand for precise, efficient proofreading, enabling more accurate reconstructions.

## 12 Acknowledgment

This work was supported by NSF award NSF-IIS-2239688 and partially supported by NIH grants 1U01NS132158 and R01HD104969. We gratefully acknowledge Amazon Web Services (AWS) for providing free cloud computing credits, which facilitated the deployment and testing of our system. We also thank the participants of our user and case studies for their valuable contributions.

